# A Computational Pipeline for Stratifying Autoimmune Patients Using Binary Antibody Data

**DOI:** 10.1101/2025.09.11.670596

**Authors:** Nathan Zhuang, Jacqueline Howells

## Abstract

This method describes a computational pipeline for stratifying autoimmune patient groups using exclusively binary autoantibody data. Our method addresses a methodological gap in computational immunology by providing a standardized framework for analyzing categorical serological data commonly found in electronic health records and resource-limited settings. The pipeline integrates three complementary analytical modules:

Module 1: Exploratory screening using statistical association tests.
Module 2: Quantification of overall immunological similarity and un-certainty.
Module 3: Prediction modeling and validation against chance.

We demonstrate the method’s utility by applying it to two autoimmune disorders. We were successful in recapitulating established clinical relationships in these two closely linked diseases. The pipeline is implemented in Python and includes detailed configuration options for custom disease groups, autoantibody panels and stratification variables. This method enables researchers to extract meaningful immunological patterns from underutilized binary clinical data, serving as a hypothesis-generation tool to help drive impactful exploration.

**Graphical Abstract:** 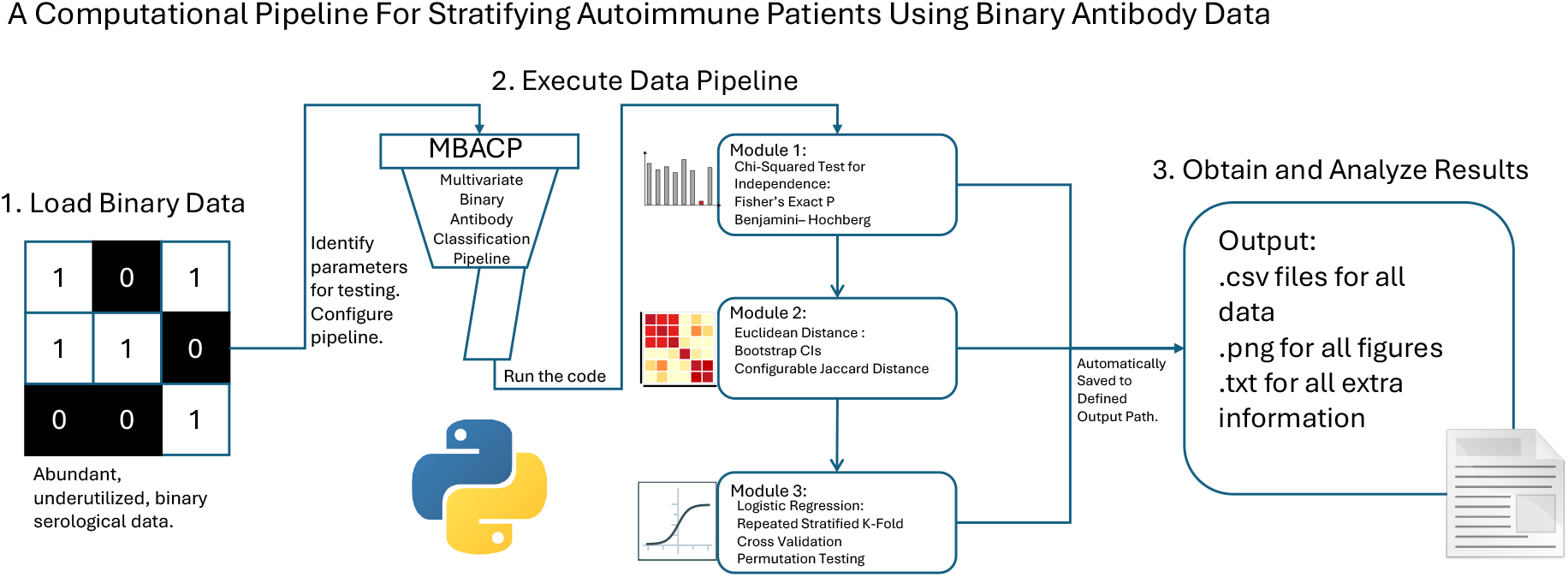

## 1. Method Name

Multivariate Binary Antibody Classification Pipeline (MBACP).

## 2. Background

### 2.1. Background and methodological gap

Binary autoantibody data represents an abundant yet underutilized resource in clinical datasets and electronic health records, particularly in resource-limited settings where continuous laboratory titers are unavailable [1, 2].Currently, no standardized computational framework exists for multivariate analysis of binary autoantibody data, forcing researchers to either use simple univariate tests, which lack power for complex pattern detection or to apply methods designed for continuous data often leading to misinterpreted results [3, 4].

Our pipeline seeks to address this gap by providing an integrated analytical framework specifically optimized for binary serological data (although continuous applications are possible). The pipeline runs statistical tests in a streamlined method that allows for easy interpretation of proposed hypotheses. The method builds upon established statistical approaches, but it combines them in a novel pipeline that maximizes statistical results and implications from categorical autoantibody data while providing validation at each analytical stage. Our pipeline can be used on all local applications that can run the latest version of Python. Virtual machines or online Python compilers will fail to read imported data as it is not localized.

## 3. Method Details

### 3.1 Pipeline architecture and workflow

The computational pipeline follows a structured workflow consisting of three main phases: data preprocessing, multi-layered statistical analysis, and comprehensive output generation. Access to all code used for dataset cleaning (if exact dataset use is desired), the pipeline (on both validated configurations), figure generation (examples in supplemental files), and datasets themselves are available in the repository listed within the specifications table.

#### 3.1.1 Data preprocessing module

The preprocessing phase handles dataset loading, cleaning, preparations for analysis, and the designation of an output file. This phase provides flexible data input capabilities that accept CSV files with customizable column names. It performs automatic data cleaning by handling missing values and creating a standard definition for each disease name. Code within the repository includes patient deduplication functionality that identifies and removes duplicate patient records based on unique identifiers and antibody profiles (not within main pipeline). Diagnosis standardization converts various diagnosis formats into standardized groups using configurable matching rules, enabling consistent group definitions across different data sources. For example, if a compiled dataset from multiple sources includes many spellings for a disease i.e. Sjogren’s and SS, they will all be standardized as a designated name within the code. Finally, the module supports subgroup stratification by clinical variables such as rheumatoid factor status, allowing for detailed subgroup analyses within broader diagnostic categories. This feature allows for targeted, high impact hypothesis generation.

#### 3.1.2 Analytical modules

The pipeline implements three complementary analytical modules that provide complementary insights into the binary autoantibody data panel selected for testing.

Module 1 performs univariate association analysis on the overall diagnostic groups (ex. SS vs RA) to assess independence between individual autoantibodies and diagnosis (i.e. if autoantibody X on its own determines diagnosis between diseases Z and Y.) This module serves as an exploratory step to verify data integrity, i.e. dataset curation, before moving onto the next modules. To do so, Chi-squared tests are implemented to assess independence between individual autoantibodies and diagnostic groups [5]. The module complements Chi-squared tests with Fisher’s exact tests, which provide exact p-values for small sample sizes or when expected cell counts are low [6]. The module also applies false discovery rate (FDR) adjustment using the Benjamini-Hochberg procedure to control multiple testing across all antibodies in the panel [7].

Module 2 conducts multivariate distance analysis to quantify overall immunological similarity between patient groups, including stratified subgroups (i.e. stratifying disease X by factor Y to see if subgroups differ). This module calculates Euclidean distances between group centroids in the multidimensional antibody space, providing a measure of overall profile similarity with correct class weightings [8]. To quantify uncertainty in these distance estimates, the module implements bootstrap confidence intervals through non-parametric resampling with 1000 iterations [9]. Additionally, the module computes Jaccard distance as an alternative similarity metric based on presence/absence patterns. However, in the repository version, the pipeline’s Jaccard distances are configured to couple all groups together and will only output 0.00. If users desire to use Jaccard distances, they must configure the metrics themselves.

Module 3 focuses on classification and validation through multivariate analysis of multiple group comparisons. Stratifying data in the same way as module 2. The module employs logistic regression with default L2 regularization (C=1.0) and balanced class weighting to prevent overfitting while maintaining interpretability and handling class imbalance [10]. Model evaluation uses repeated stratified k-fold cross-validation with different iteration schemes: main performance metrics (accuracy, balanced accuracy, ROC AUC) employ 5 folds with 10 repeats (50 total iterations) for robust estimation, while ROC curve generation uses 5 folds with 2 repeats (10 iterations) [11]. Current iteration numbers are all customizable to any value desired, the two different iterations are used in the default configuration to show the capability of the pipeline. Statistical significance of classification performance is assessed through permutation testing with 100 label randomizations (again, customizable), each evaluated using 5-fold cross-validation with 2 repeats, ensuring that observed performance exceeds chance levels [12]. The module reports comprehensive performance metrics including accuracy, balanced accuracy, and ROC AUC with appropriate uncertainty estimates [13].

#### 3.1.3. Output generation

The pipeline generates output designed to support both methodological validation and scientific interpretation (hypothesis generation). These outputs include statistical summary tables in CSV, visualizations such as confusion matrices and bootstrap distributions to illustrate key findings, and complete reproducibility information including code hashes and package versions.

### 3.2. Software requirements and installation

The pipeline requires Python 3.9 or higher with several essential scientific computing packages. The core dependencies include pandas (*≥*1.4.0) for data manipulation, numpy (≥1.21.0) for numerical computations, scikit-learn (≥1.0.0) for machine learning algorithms, scipy (≥1.7.0) for statistical functions, matplotlib (≥3.5.0) and seaborn (≥0.11.0) for visualization, and statsmodels (≥0.13.0) for advanced statistical testing. Installation is straightforward as the pipeline consists of a single Python script with no complex dependencies or external compilation requirements. All package versions used to create the results communicated in this paper are available in the README.txt file within the repository.

### 3.3 Data preparation and formatting

#### 3.3.1 Required data structure

The pipeline expects input data in CSV format with a specific structure or errors will occur. The data should be organized with patients as rows and variables as columns, where the columns represent the autoantibodies being tested (does not have to be the exact panel, the preprocessing platform will sort out only the columns wanted). Table 1 outlines the required columns and their specifications. The data can a unique patient identifier column but must include: a diagnosis column containing disease labels, one stratification variable encoded as binary values, and 1 or more autoantibody columns representing the “autoantibody panel”, all encoded as binary (0/1) variables. However, datasets with titers can also be processed with this pipeline if titers are converted to binary values by some metric (i.e. abnormal levels vs healthy levels can be coded in a binary format.)

**Table 1:**
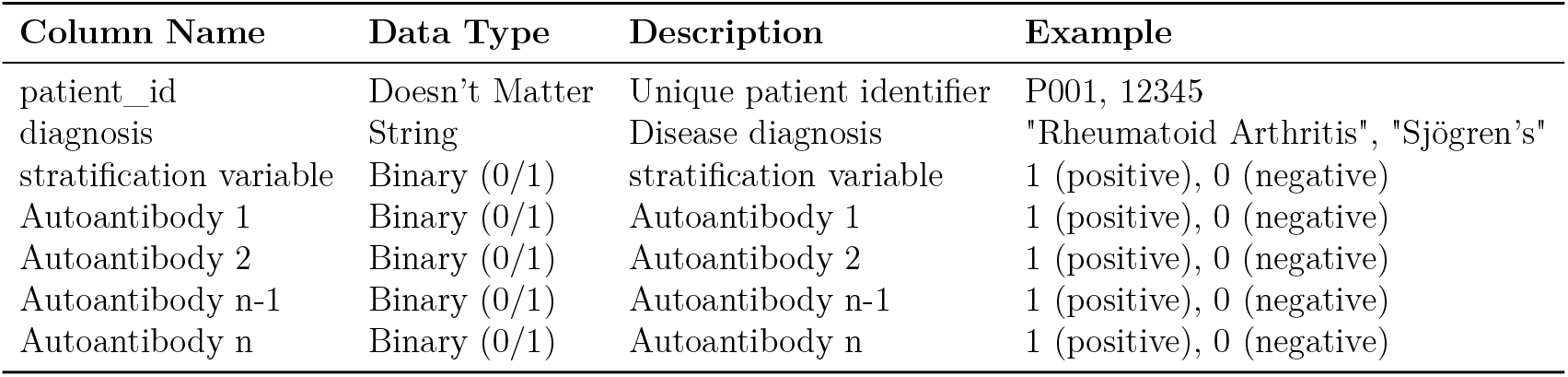
Required data structure for pipeline input.

#### 3.3.2 Customizing antibody panels

Users can easily modify the autoantibody panel to match their specific research needs by editing the main code. The default configuration includes four common autoantibodies (used in validation), which can be customized as shown in the following code example:

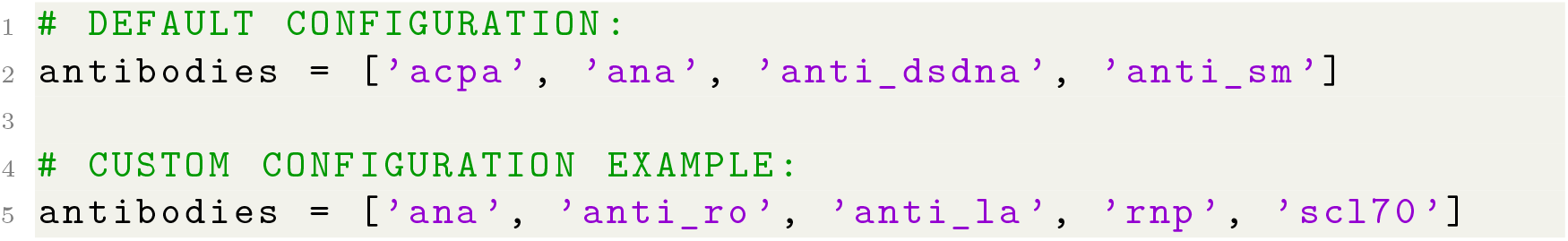

The pipeline automatically adapts to the specified antibody columns, provided they exist in the input data and contain properly encoded binary values. The antibody names must match the column names in the CSV file exactly. If the column does not match, or the column is not present within the dataset, the code will run without that specified autoantibody and the console will print the column as not included.

#### 3.3.3 Customizing disease groups

Disease group definitions can be modified in the `standardize_diagnosis` function to accommodate different spelling and formatting if using a compiled dataset. The function uses regular expressions to map various diagnosis formats to standardized group labels, as demonstrated in this code implementation:

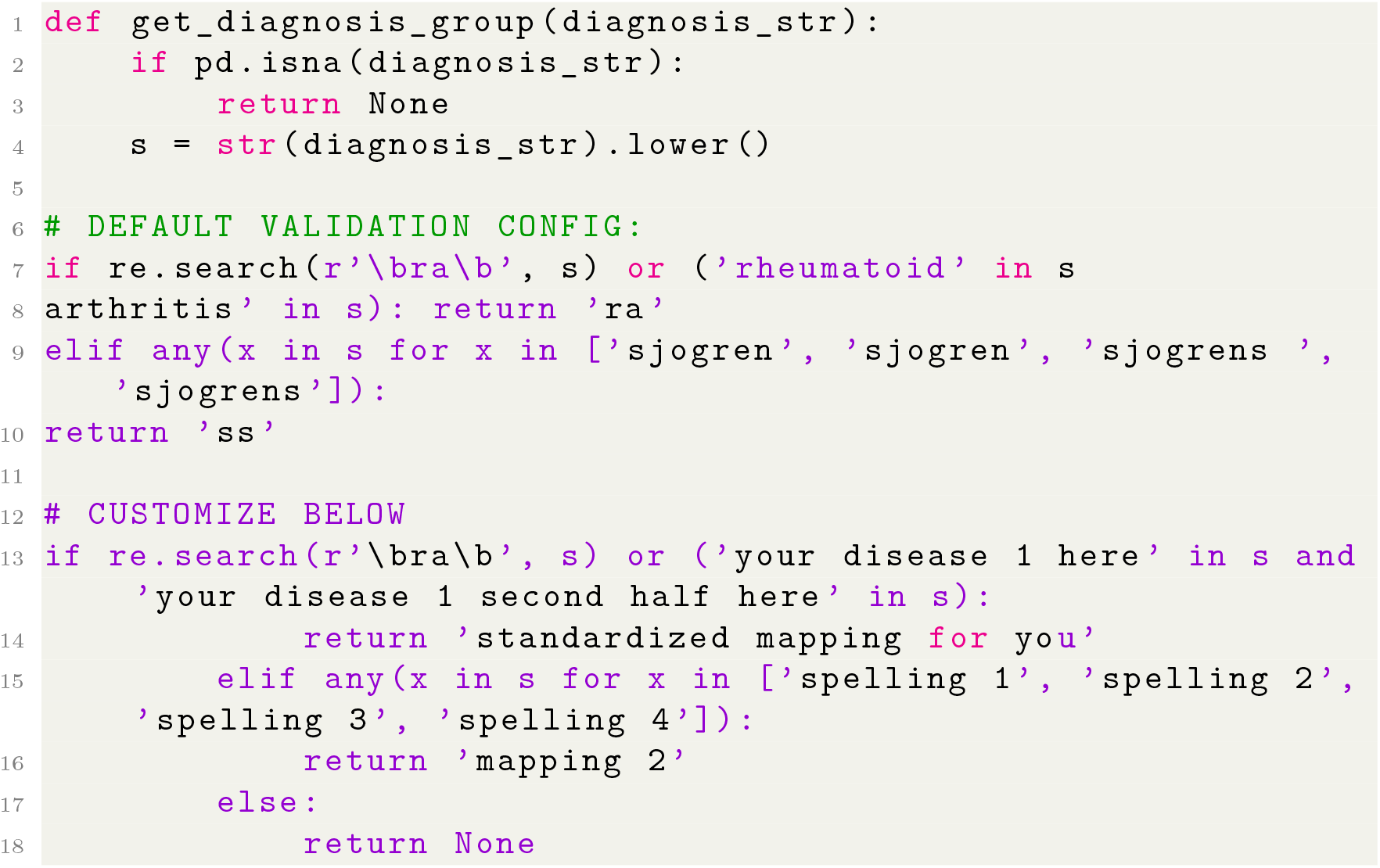

Researchers can extend the existing mapping by adding new disease patterns or modify the existing patterns to better match their specific dataset terminology. This flexibility allows the pipeline to be applied to diverse autoimmune disease research questions beyond the original RA and SS focus.

#### 3.3.4 Customizing stratification variables

The pipeline supports stratification by any binary clinical variable present in the dataset, enabling detailed subgroup analyses. The default implementation uses rheumatoid factor status to stratify Sjögren’s syndrome patients, but researchers can easily modify this to use other clinically relevant variables. Stratification is defined separately in two functions: `run_logistic_analysis` and `distance_analysis`, as shown in this code example:

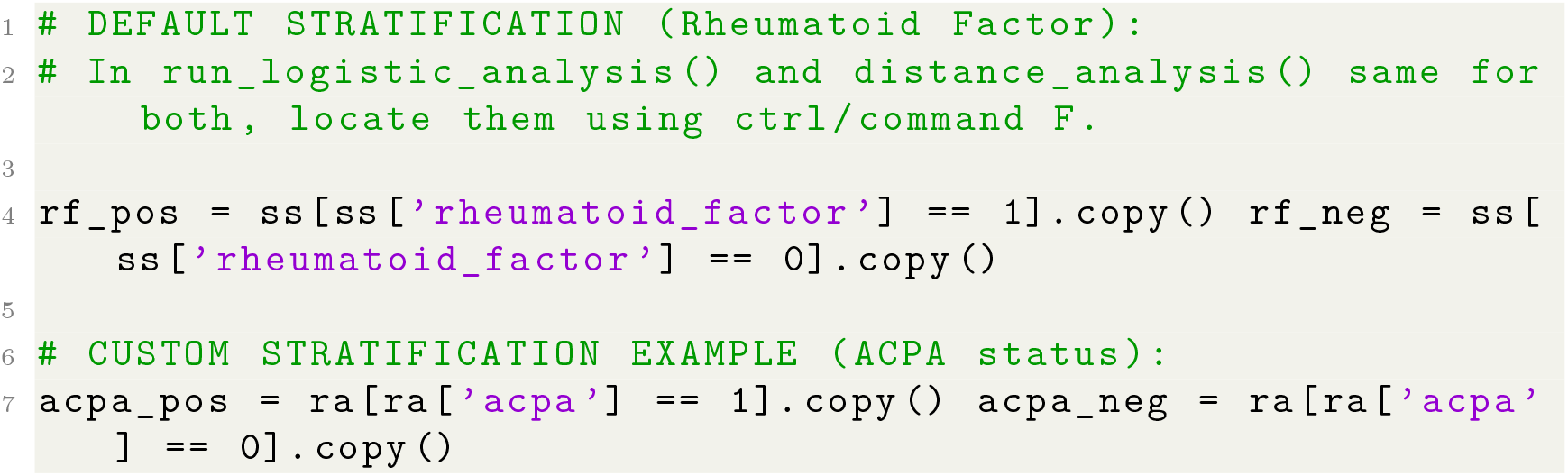

The stratification logic is implemented in clearly marked sections of the analysis functions, making customization straightforward for users with basic Python knowledge. Any binary column in the dataset can be used for stratification purposes.

### 3.4 Running the analysis

#### 3.4.1. Basic execution

The pipeline can be executed with default settings using a simple command line invocation:

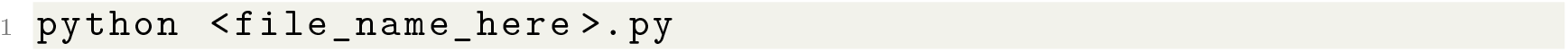

Running the script automatically triggers the complete analytical workflow. The default configuration includes disease and autoantibody profiles that have been validated on two datasets.

#### 3.4.2. Configuration options

Several key configuration parameters are hardcoded within specific functions and can be modified to adapt the pipeline to specific research needs. Users can adjust these parameters directly in the source code:

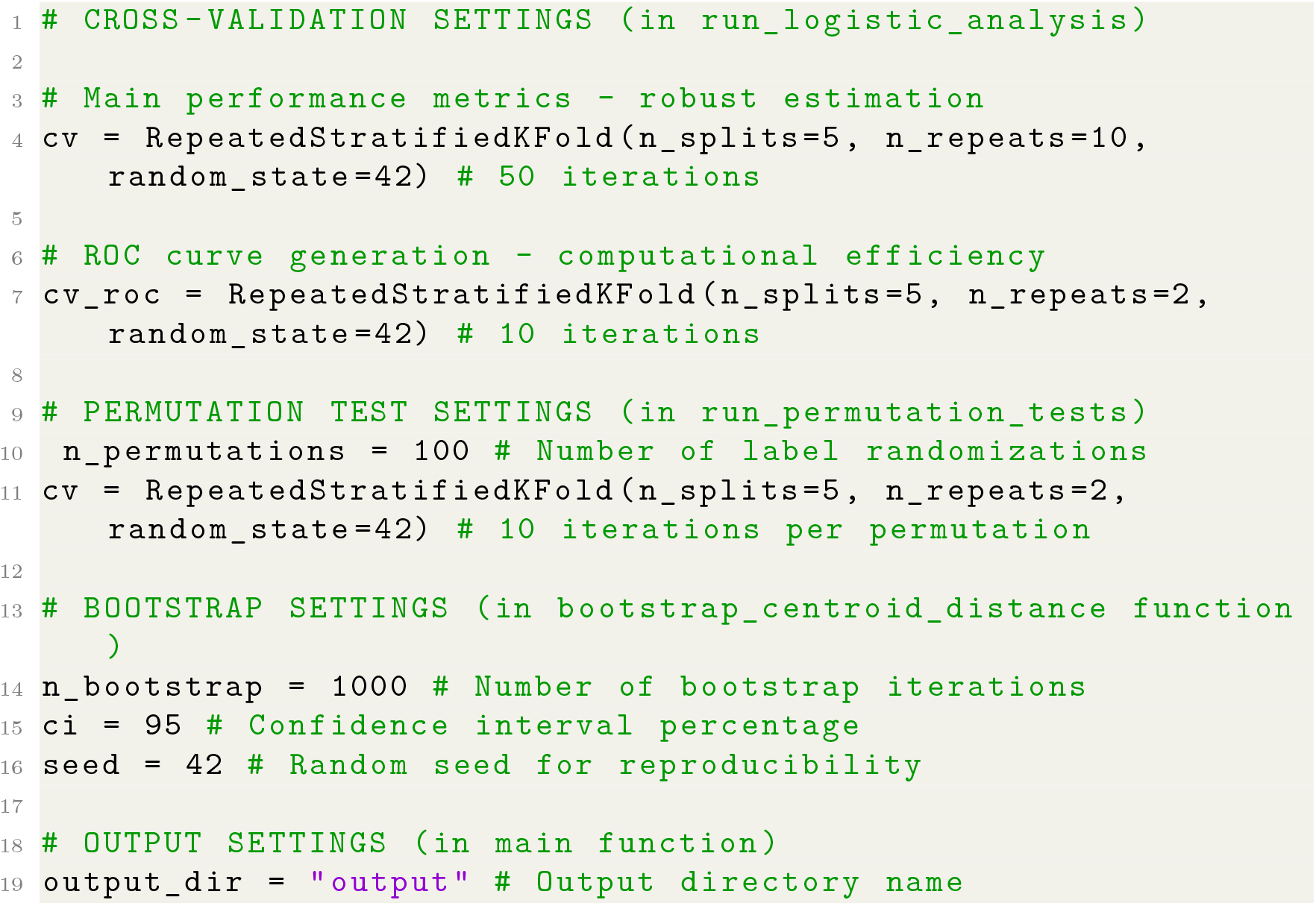

Cross-validation settings control the robustness of performance estimation, with main metrics using 50 iterations (5 folds × 10 repeats) for stable estimates, while ROC curves and permutation tests use 10 iterations (5 folds × 2 repeats) for computational efficiency. Permutation testing performs 100 label randomizations, each evaluated with 10-fold cross-validation. Bootstrap resampling uses 1000 iterations with 95% confidence intervals by default. Output settings allow researchers to specify custom directory names for organizing results. By upping each value, the time and memory strain on the performer’s device will increase. The current values run smoothly on most laptops on decently sized datasets but have not been tested on any datasets with N>1500 or increased values of any of these configurations.

#### 3.4.3. Handling special cases

The pipeline includes specific features to handle common data challenges encountered in real-world research. For datasets with severe class imbalance, the pipeline automatically adjusts class weights inversely proportional to class frequencies using the `class_weight=‘balanced’` parameter in logistic regression. This ensures equal importance of minority and majority classes during model training:

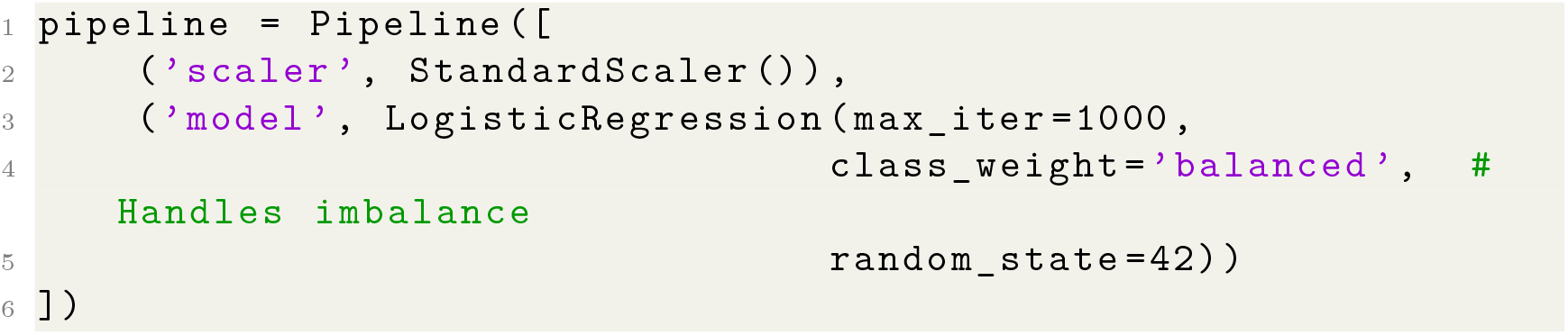

When working with small datasets, users may benefit from increasing the number of permutation tests for more precise p-value estimation and should rely more heavily on Fisher’s exact tests in the univariate analysis, which the pipeline automatically computes alongside Chi-squared tests.

### 3.5. Output interpretation

#### 3.5.1. Key output files

The pipeline generates a comprehensive set of output files designed to support complete methodological documentation and result interpretation. The `classifier_performance.csv` file contains cross-validated performance metrics for all group comparisons, including accuracy, balanced accuracy, and ROC AUC with standard deviations (based on 50 CV iterations). The `statistical_analysis.csv` file provides results of univariate association tests for each antibody comparing overall SS vs RA groups, including Chisquared and Fisher’s exact test statistics, p-values, adjusted p-values, and odds ratios. Distance analysis results are saved in `distance_analysis.csv` containing Euclidean and Jaccard distance metrics with bootstrap confidence intervals for multiple group comparisons including RF-stratified subgroups. Raw 2×2 contingency tables for each antibody are exported in `contingency_tables.csv` for additional verification. Reproducibility information including code hashes, timestamps, and package versions is documented in `version_info.txt`. Additionally, the pipeline generates multiple visualization plots including ROC curves with confidence intervals, bootstrap distribution histograms, confusion matrices, and permutation test results for intuitive result interpretation. All results are also saved in `all_results.pkl` as a Python pickle file for programmatic access.

#### 3.5.2. Interpreting distance metrics

To interpret the results that the pipeline generates, a basic understanding of p-values and thresholds is needed:

For chi-squared tests in module 1, a p value greater than 0.05 signals that disease diagnosis is not dependent on the autoantibody [5]. For example, if a p value of 0.1 was determined for autoantibody X in stratifying diseases X and Y, then the disease diagnosis is not dependent on the positivity status of autoantibody X. Fisher’s OR and Benjami-Hochberg FDR both accompany the chi squared tests to account for small sample size and false positives respectively. Fisher’s can be interpreted in the same way as chi-squared p-values [6]. While the Benjamini-Hochberg procedure does not change the test itself; rather, it adjusts the p-value to account for the number of comparisons and control the FDR. Adjusted p-values are then compared to the significance threshold in the same way as ordinary p-values but reflect FDR control [7].

Euclidean distances range from 0, indicating identical antibody profiles between groups, to 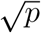 (where *p* is the number of antibodies) in extreme cases of completely dissimilar profiles. Smaller distance values suggest greater immunological similarity [8]. The bootstrap confidence intervals accompanying each distance estimate provide important information about measurement precision, with wider intervals typically indicating greater uncertainty often associated with smaller sample sizes [9].

Classification performance metrics offer insights into the discriminative power of antibody profiles for distinguishing between patient groups. ROC AUC values close to 0.5 indicate classification performance indistinguishable from random guessing, while values approaching 1.0 suggest strong discriminative ability. Values between 0.7-0.8 are often considered acceptable discrimination, 0.8-0.9 indicate excellent discrimination, and above 0.9 suggest outstanding discrimination [10, 12]. Permutation p-values below 0.05 indicate that the observed classification performance significantly exceeds chance levels, providing statistical validation of the multivariate patterns [13]. These p-values are calculated by comparing the true ROC AUC (from 10 CV iterations) against the distribution of 100 permuted label AUCs. Confidence intervals around performance metrics (expressed as standard deviations from 50 CV iterations) help assess estimate stability, with smaller standard deviations generally reflecting more reliable performance estimates.

The relationship between distance metrics and classification performance often shows an inverse pattern, where immunologically distinct groups (larger Euclidean distances) are typically easier to classify (higher AUC values), providing internal validation of the analytical consistency.

## 4. Method validation

We validated the pipeline using both synthetic and clinical autoimmune datasets to assess performance under controlled and real-world conditions.

### 4.1. Validation datasets

The synthetic validation used the Comprehensive Autoimmune Disorder Dataset [14], which contained 176 patients after deduplication, including 91 rheumatoid arthritis (RA) patients and 85 Sjögren’s syndrome (SS) patients with rheumatoid factor stratification (RF-positive SS: n=38, RF-negative SS: n=47). For real-world validation, we applied the pipeline to the Diagnosis of Rheumatic and Autoimmune Diseases dataset [15], which included 1,229 patients (RA: n=766, SS: n=463) with substantial class imbalance in RF stratification among SS patients (RF-positive SS: n=433, RF-negative SS: n=30). In the clinical dataset, many autoantibodies were used as diagnostic specifiers, meaning 100% positive as it was required for a diagnosis of that disease. This led to a constraint in selecting an antibody profile. We settled with HLA-B27, anti-dsDNA and anti-Sm. Each has a overall autoimmune impact that we hoped would produce meaningful relationships.

### 4.2. Statistical analysis

The pipeline successfully performed comprehensive statistical analyses on both datasets. For the synthetic dataset, univariate association tests between RA and SS groups revealed no statistically significant associations after Benjamin-Hochberg correction. ANA showed the strongest trend toward association (*χ*^2^=2.910, p=0.0880, adjusted p=0.3242; Fisher’s OR=0.531, p=0.0689, adjusted p=0.2651), while other antibodies showed minimal association: ACPA (*χ*^2^=0.005, p=0.9419; Fisher’s OR=0.933, p=0.8790), anti-dsDNA (*χ*^2^=1.954, p=0.1621; Fisher’s OR=0.624, p=0.1325), and anti-Sm (*χ*^2^=0.167, p=0.6830; Fisher’s OR=1.185, p=0.6497). In the clinical dataset, similar non-significant patterns emerged for the available antibody panel: HLA-B27 (*χ*^2^=0.000, p=0.9823; Fisher’s OR=1.010, p=0.9531), anti-dsDNA (*χ*^2^=0.201, p=0.6536; Fisher’s OR=0.942, p=0.6378), and anti-Sm (*χ*^2^=0.657, p=0.4176; Fisher’s OR=1.108, p=0.4091). All clinical dataset p-values remained non-significant after multiple testing correction.

Euclidean distance analysis with bootstrap confidence intervals revealed coherent immunological similarity patterns that aligned with established clinical knowledge in both datasets (Table **??**). In the synthetic dataset, RF-positive SS patients showed significantly closer immunological proximity to RA (distance = 0.195, 95% CI: 0.068-0.335) compared to RF-negative SS patients (distance = 0.348, 95% CI: 0.197-0.507). The RF-positive SS vs RF-negative SS comparison showed the largest distance (0.362, 95% CI: 0.173-0.558), while the overall SS vs RA distance was intermediate (0.223, 95% CI: 0.095-0.362).

**Figure 1:**
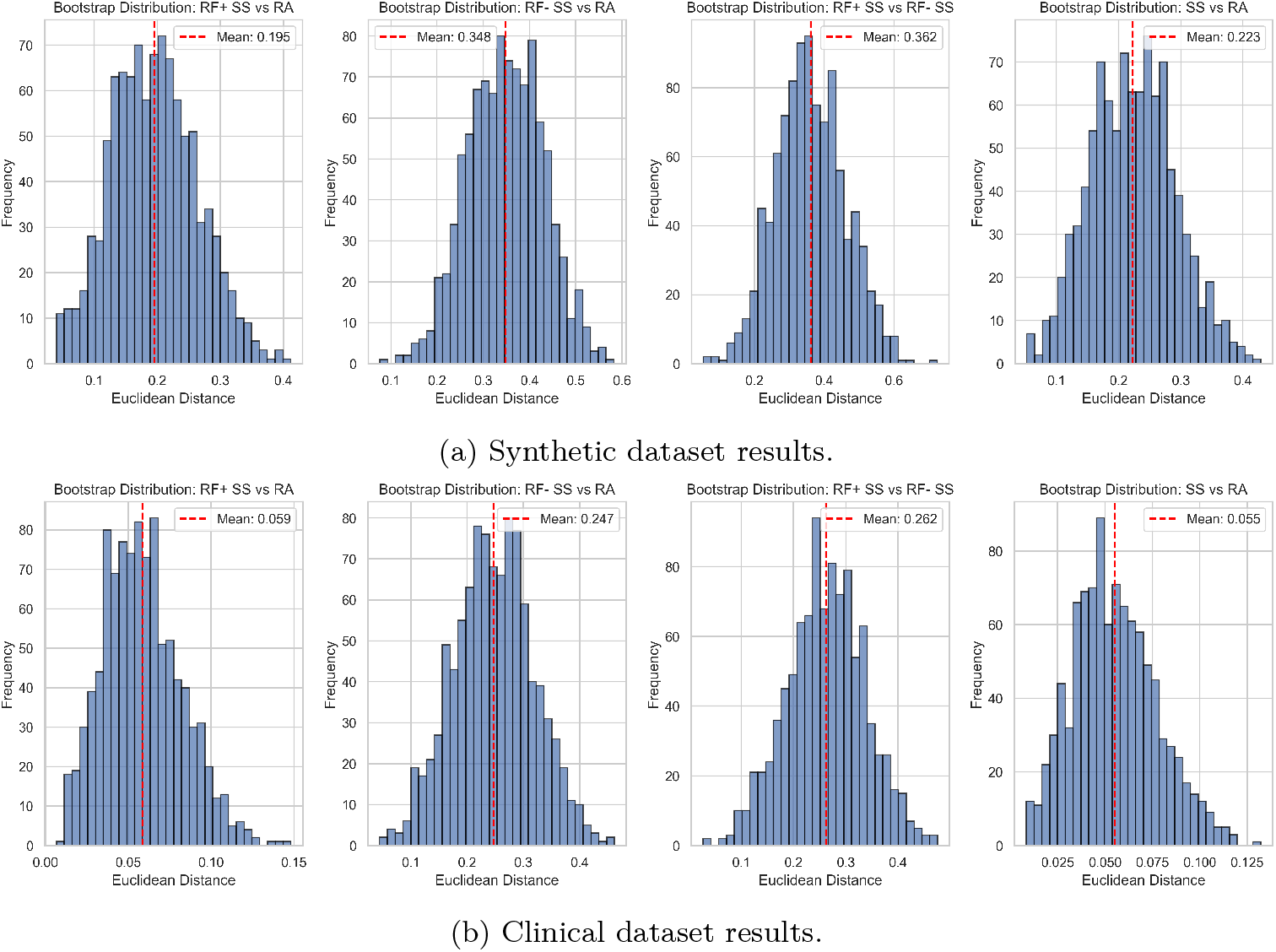
Bootstrap analyses of immunological distance metrics. Empirical Euclidean-distance distributions from 1000 bootstrap iterations. Panel (a) shows results obtained from the synthetic dataset, while panel (b) shows corresponding distributions from the clinical dataset. This is one example output from the pipeline included to show the pipeline’s visual capabilities.

The clinical dataset reproduced this directional pattern despite different antibody panels and substantial class imbalance. RF-positive SS patients showed closer serological proximity to RA (distance = 0.059, 95% CI: 0.019-0.108) compared to RF-negative SS patients (distance = 0.247, 95% CI: 0.108-0.389). The consistency across datasets demonstrates the pipeline’s ability to capture meaningful biological signals to help aid hypothesis generation even in different data characteristics.

Logistic regression models demonstrated performance patterns that directly reflected the immunological relationships identified through distance analysis (Table 2). In the synthetic dataset, the RF-negative SS vs RA comparison showed the highest discriminative ability (ROC AUC = 0.639 ± 0.093), consistent with their greater immunological distance. Conversely, RF-positive SS vs RA showed minimal predictive power (ROC AUC = 0.362 ± 0.102), accurately reflecting their serological similarity. The RF-positive SS vs RF-negative SS comparison showed intermediate performance (ROC AUC = 0.618 ± 0.130), while overall SS vs RA discrimination was modest (ROC AUC = 0.557 ± 0.088).

**Table 2:** Classification Performance with Permutation Test Results.

| Comparison | Accuracy | Balanced Accuracy | ROC AUC | Permutation p | Class Ratio |
| --- | --- | --- | --- | --- | --- |
| **Synthetic Dataset** |  |  |  |  |  |
| RF-positive SS vs RA | $0.391 \pm 0.091$ | $0.364 \pm 0.080$ | $0.362 \pm 0.102$ | 0.9406 | 1:2.4 |
| RF-positive vs RF-negative SS | $0.608 \pm 0.113$ | $0.602 \pm 0.115$ | $0.618 \pm 0.130$ | 0.0594 | 1:1.2 |
| RF-negative SS vs RA | $0.635 \pm 0.073$ | $0.639 \pm 0.073$ | $0.639 \pm 0.093$ | 0.0396 | 1:1.9 |
| SS vs RA | $0.566 \pm 0.085$ | $0.568 \pm 0.085$ | $0.557 \pm 0.088$ | 0.1881 | 1:1.1 |
| **Clinical Dataset** |  |  |  |  |  |
| RF-positive SS vs RA | $0.485 \pm 0.028$ | $0.485 \pm 0.027$ | $0.490 \pm 0.033$ | 0.4752 | 1:1.8 |
| RF-positive vs RF-negative SS | $0.591 \pm 0.085$ | $0.547 \pm 0.097$ | $0.580 \pm 0.102$ | 0.0891 | 14.4:1 |
| RF-negative SS vs RA | $0.553 \pm 0.071$ | $0.547 \pm 0.087$ | $0.572 \pm 0.081$ | 0.0693 | 1:25.5 |
| SS vs RA | $0.486 \pm 0.029$ | $0.486 \pm 0.024$ | $0.482 \pm 0.026$ | 0.8020 | 1:1.7 |

In the clinical dataset, classification performance was substantially impacted by class imbalance despite class-weighted logistic regression. RF-positive SS vs RA achieved near-chance performance (ROC AUC = 0.490 ± 0.033, permutation p = 0.4752), while RF-negative SS vs RA showed modest but non-significant discrimination (ROC AUC = 0.572 ± 0.081, permutation p = 0.0693). The extreme class imbalance in RF-positive vs RF-negative SS comparison (14.4:1 ratio) resulted in challenging classification (ROC AUC = 0.580 ± 0.102, permutation p = 0.0891). Permutation testing provided crucial statistical context for classification performance across both datasets. In the synthetic dataset, only the RF-negative SS vs RA comparison achieved statistical significance (p = 0.0396), while the RF-positive SS vs RA comparison showed performance indistinguishable from chance (p = 0.9406). The RF-positive SS vs RF-negative SS comparison approached but did not reach significance (p = 0.0594). In the clinical dataset, no comparisons reached statistical significance, with all permutation p-values exceeding 0.05, reflecting the greater analytical challenge posed by real-world data characteristics.

Analysis of autoantibody prevalence revealed distinct serological profiles across diagnostic groups in both datasets. In the synthetic dataset, RF-negative SS patients demonstrated higher prevalence of ANA (83.0% vs 71.1%) and anti-dsDNA (72.3% vs 44.7%) compared to RF-positive SS patients, while ACPA and anti-Sm prevalence remained similar across sub-groups. In the clinical dataset, RF-negative SS patients exhibited higher HLA-B27 prevalence (63.3% vs 49.0%) but lower anti-dsDNA prevalence (36.7% vs 52.0%) compared to RF-positive SS patients. These patterns demonstrate the pipeline’s ability to capture clinically relevant serological heterogeneity across different antibody panels and population characteristics.

Consistent with our findings, RF-positive Sjögren’s syndrome patients exhibit immunological features more like rheumatoid arthritis than RF-negative patients. Specifically, RF binding profiles in RF-positive SS closely resemble those in RA, whereas RF-negative SS shows distinct patterns [16]. This supports the validity of using RF status to define an RA-like immunological subtype within SS.

## 5. Limitations

This method will not work if biomarkers (e.g., antibodies) selected for stratification are also diagnostic or inclusion criteria within the specific dataset being analyzed. For instance, if Antibody M is used as a diagnostic specifier for Disease X during the original data collection, then all subjects classified as having Disease X in the dataset will be uniformly positive for Antibody M. In this scenario, attempting to use Antibody M to stratify Disease X methodologically unsound. Furthermore, the inclusion of such a diagnostically specified marker (e.g., Antibody M) anywhere within the broader panel of biomarkers used in the analysis pipeline will introduce confounding noise and reduce the power of the method. Even if stratification is based on a separate factor (e.g., Antibody N or Factor Y), the uniform positivity of all Disease X cases for Antibody M means this marker will appear highly predictive of Disease X, effectively creating a false signal or artifact that disrupts the intended stratification outcome. This scenario results in an uninformative analysis, as the artifactual signal obscures the true biological differences the method is designed to uncover. Researchers should therefore ensure that the factors used for stratification are independent of the diagnostic criteria used to generate the dataset under investigation.

The method’s performance in terms of memory usage and execution time is primarily dictated by the size of the input data and the chosen parameters for statistical validation. The calculation of bootstrap confidence intervals (CIs) and the execution of permutation tests are computationally demanding, particularly regarding memory allocation on minimal operating systems. This cost is directly proportional to the number of iterations specified for both tests. To manage computational resources, users can easily configure this. Execution time scales positively with the size of the input data. Specifically, processing time increases with both the number of subjects in the dataset and the number of biomarkers (e.g., antibodies) included in the panel.

While the MBACP is optimized for binary input, we recognize that when continuous laboratory measurements are converted to binary values (dichotomization), this process results in a loss of statistical power and information relative to methods that utilize the full range of continuous values.

## Supporting information

All outputs from pipeline - Supplemental

## Ethics Statement

This method uses de-identified data; no ethical approval was required for the development and validation of the computational pipeline. Users should ensure appropriate ethical approvals for their specific datasets and research contexts.

## CRediT Authorship Contribution Statement

**Nathan Zhuang**: Methodology, Software, Validation, Formal analysis, Data curation, Writing - original draft, Visualization.

**Jacqueline Howells**: Supervision, Writing - review & editing, Project administration.

## Declaration of Competing Interests

The authors declare no competing interests.

## Acknowledgments

The authors thank Polygence for providing a platform and support to conduct this research. The method development was supported by the Polygence mentorship program.

## Data and Code Availability

The complete pipeline code, configuration files, and usage examples are available at: [GitHub Repository](https://github.com/NathanZhuangCalgary/Multivariate-Binary-Antibody-Classification-Pipeline-MBACP)

## Notes

https://doi.org/10.34740/kaggle/dsv/9807734

https://github.com/NathanZhuangCalgary/Multivariate-Binary-Antibody-Classification-Pipeline-MBACP

https://doi.org/10.1016/j.dib.2025.111623

### Competing Interest Statement

This work was developed as part of the Polygence mentorship program. Polygence provided guidance on research design but had no role in data collection/analysis or manuscript.

### Summary of Updates

Overall revisions to grammar; limitations updated.

