## Supplementary material for "A Computational Pipeline for Stratifying Autoimmune Patients Using Binary Antibody Data": All outputs from pipeline - Supplemental

---

---

### 1. Supplementary Materials Overview

This file contains all outputs generated by the pipeline, all figures and tables are represented across both validations preformed within the article. Additional figure generation scripts are present within the linked repository, but they are only for easier interpretation and do not affect the results obtained in any way. The first section displays results obtained from the synthetic dataset, and the second section displays the results obtained from clinical validation. As a whole, these supplementary materials are presented to show the full capability of the pipeline, including all results and figures. Jaccard distances were omitted from the results and calibrated to show 0.00 in the current pipline, results reflect this.

### 2. Synthetic Data Results

#### *2.1. Figure Outputs*

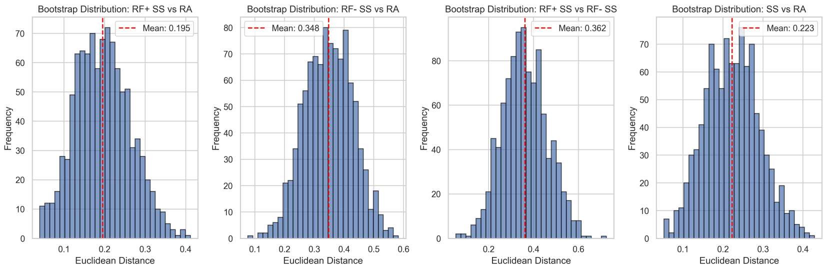


Figure 1: Bootstrap distributions showing the variability in performance metrics across different classifier comparisons using synthetic data. The distributions provide insight into the stability and reliability of the classification results.

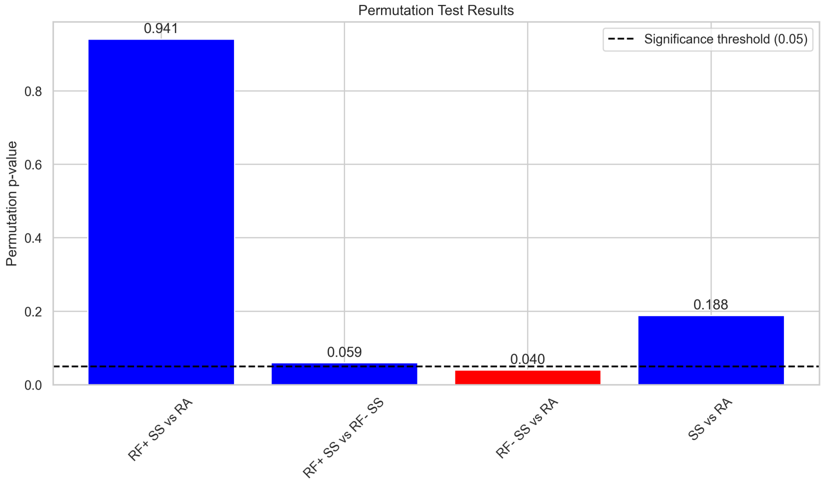


Figure 2: Permutation test results for different classifier comparisons using synthetic data. The dashed line indicates the significance threshold (0.05). Results show permutation p-values of 0.059 for RF+ SS vs RA, 0.040 for RF+ SS vs RF- SS, and 0.188 for RF- SS vs RA.

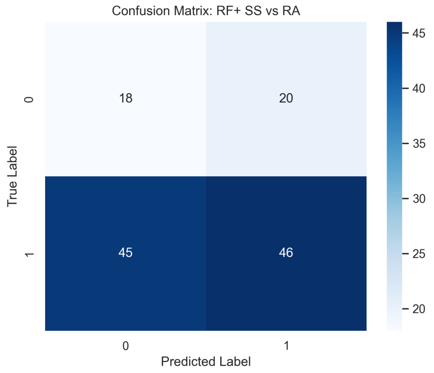


(a) RF+ SS vs RA

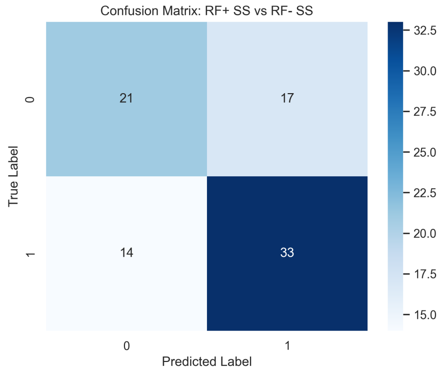


(b) RF+ SS vs RF- SS

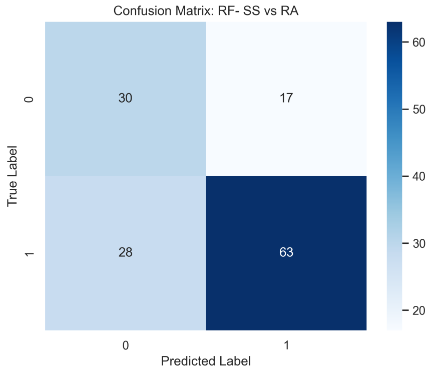


(c) RF- SS vs RA

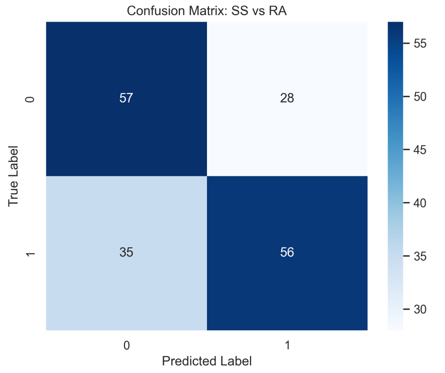


(d) SS vs RA

Figure 3: Confusion matrices for different classifier comparisons using synthetic data showing true vs predicted labels. The matrices provide detailed insight into classification performance across different experimental conditions.

#### *2.2. Table Outputs*

Table 1: Classifier Performance Metrics using Synthetic Data (Mean ± Standard Deviation)

| **Comparison** | **Accuracy** | **Balanced Accuracy** | **ROC AUC** |
|---|---|---|---|
| RF+ SS vs RA | 0.391 ± 0.091 | 0.364 ± 0.080 | 0.362 ± 0.102 |
| RF+ SS vs RF- SS | 0.608 ± 0.113 | 0.602 ± 0.115 | 0.618 ± 0.130 |
| RF- SS vs RA | 0.635 ± 0.073 | 0.639 ± 0.073 | 0.639 ± 0.093 |
| SS vs RA | 0.566 ± 0.085 | 0.568 ± 0.085 | 0.557 ± 0.088 |

Table 2: Statistical Analysis of Antibody Associations using Synthetic Data

| Antibody | Chi-square | p-value (Chi-square) | p-value (Fisher) | Odds Ratio | Chi-square Valid | Min Expected |
|---|---|---|---|---|---|---|
| acpa | 0.00532 | 0.94187 | 0.87902 | 0.93333 | True | 35.74 |
| ana | 2.91037 | 0.08801 | 0.06894 | 0.53078 | True | 24.63 |
| anti_dsdna | 1.95445 | 0.16211 | 0.13254 | 0.62411 | True | 39.12 |
| anti_sm | 0.16678 | 0.68299 | 0.64965 | 1.18519 | True | 38.15 |

Table 3: Contingency Tables for Antibody Analysis using Synthetic Data

| Group | Positive | Negative |
|---|---|---|
| RA_acpa | 52 | 39 |
| SS_acpa | 50 | 35 |
| RA_ana | 59 | 32 |
| SS_ana | 66 | 19 |
| RA_anti_dsdna | 44 | 47 |
| SS_anti_dsdna | 51 | 34 |
| RA_anti_sm | 52 | 39 |
| SS_anti_sm | 45 | 40 |

Table 4: Distance Analysis Between Groups using Synthetic Data

| Comparison | Euclidean Distance | Euclidean CI Lower | Euclidean CI Upper | Jaccard Distance | n1 | n2 |
|---|---|---|---|---|---|---|
| RF+ SS vs RA | 0.19533 | 0.06838 | 0.33535 | 0.0 | 38 | 91 |
| RF- SS vs RA | 0.34825 | 0.19735 | 0.50717 | 0.0 | 47 | 91 |
| RF+ SS vs RF- SS | 0.36165 | 0.17294 | 0.55843 | 0.0 | 38 | 47 |
| SS vs RA | 0.22293 | 0.09537 | 0.36150 | 0.0 | 85 | 91 |

Table 5: Version Information for Synthetic Data Analysis

| Component | Version/Value |
|---|---|
| Code Hash | a3c124391aa36169 |
| Timestamp | 2025-11-11 10:56:14 |
| Python Version | 3.13.3 |
| Pandas | 2.2.3 |
| NumPy | 2.2.5 |
| Scikit-learn | 1.7.2 |
| SciPy | 1.16.2 |

### 3. Clinical Data Results

#### *3.1. Figure Outputs*

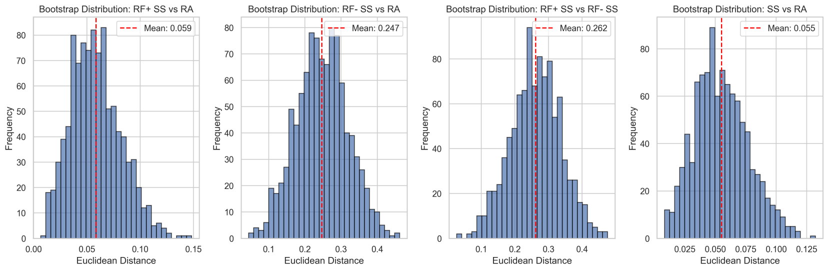


Figure 4: Bootstrap distributions showing the variability in performance metrics across different classifier comparisons using clinical data. The distributions provide insight into the stability and reliability of the classification results.

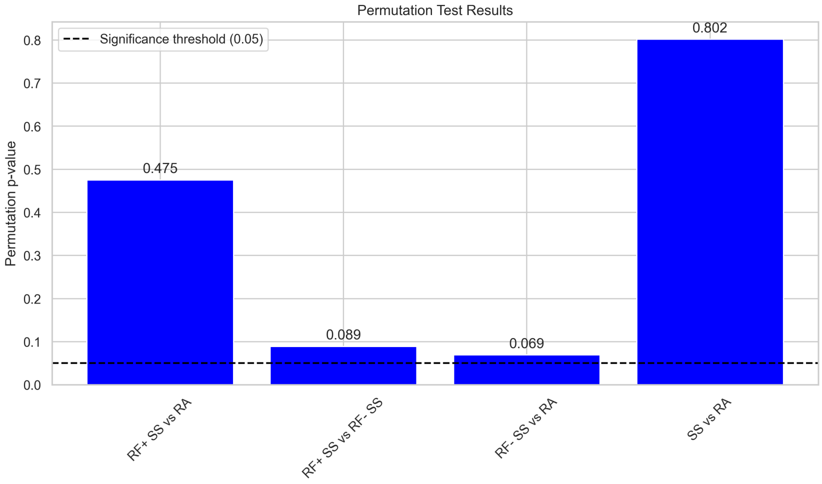


Figure 5: Permutation test results for different classifier comparisons using clinical data. The dashed line indicates the significance threshold (0.05). Results show the statistical significance of performance differences between classifiers.

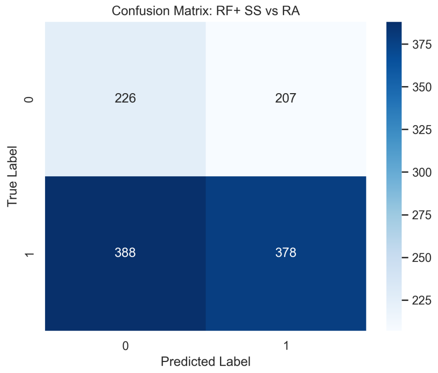


(a) RF+ SS vs RA

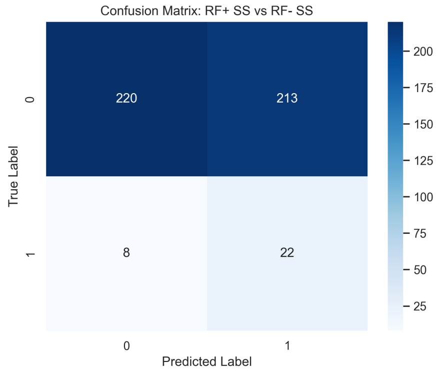


(b) RF+ SS vs RF- SS

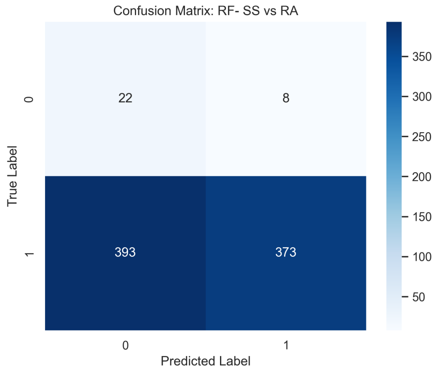


(c) RF- SS vs RA

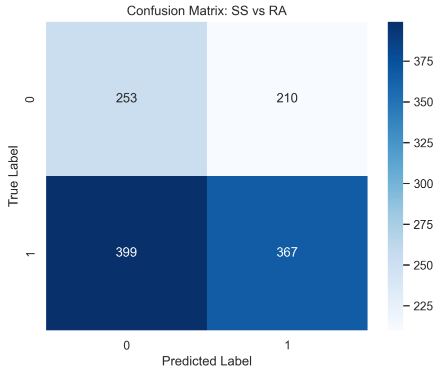


(d) SS vs RA

Figure 6: Confusion matrices for different classifier comparisons using clinical data showing true vs predicted labels. The matrices provide detailed insight into classification performance across different experimental conditions.

#### *3.2. Table Outputs*

Table 6: Classifier Performance Metrics using Clinical Data (Mean ± Standard Deviation)

| **Comparison** | **Accuracy** | **Balanced Accuracy** | **ROC AUC** |
|---|---|---|---|
| RF+ SS vs RA | 0.485 ± 0.028 | 0.485 ± 0.027 | 0.490 ± 0.033 |
| RF+ SS vs RF- SS | 0.591 ± 0.085 | 0.547 ± 0.097 | 0.580 ± 0.102 |
| RF- SS vs RA | 0.553 ± 0.071 | 0.547 ± 0.087 | 0.572 ± 0.081 |
| SS vs RA | 0.486 ± 0.029 | 0.486 ± 0.024 | 0.482 ± 0.026 |

Table 7: Statistical Analysis of Antibody Associations using Clinical Data

| Antibody | Chi-square | p-value (Chi-square) | p-value (Fisher) | Odds Ratio | Chi-square Valid | Min Expected |
|---|---|---|---|---|---|---|
| hla_b27 | 0.00049 | 0.98231 | 0.95309 | 1.00959 | True | 231.31 |
| anti_dsdna | 0.20138 | 0.65361 | 0.63779 | 0.94198 | True | 231.31 |
| anti_sm | 0.65716 | 0.41756 | 0.40906 | 1.10814 | True | 217.37 |

Table 8: Contingency Tables for Antibody Analysis using Clinical Data

| Group | Positive | Negative |
|---|---|---|
| RA_hla_b27 | 384 | 382 |
| SS_hla_b27 | 231 | 232 |
| RA_anti_dsdna | 379 | 387 |
| SS_anti_dsdna | 236 | 227 |
| RA_anti_sm | 367 | 399 |
| SS_anti_sm | 210 | 253 |

Table 9: Distance Analysis Between Groups using Clinical Data

| Comparison | Euclidean Distance | Euclidean CI Lower | Euclidean CI Upper | Jaccard Distance | n1 | n2 |
|---|---|---|---|---|---|---|
| RF+ SS vs RA | 0.05853 | 0.01891 | 0.10791 | 0.0 | 433 | 766 |
| RF- SS vs RA | 0.24699 | 0.10786 | 0.38868 | 0.0 | 30 | 766 |
| RF+ SS vs RF- SS | 0.26175 | 0.11201 | 0.40891 | 0.0 | 433 | 30 |
| SS vs RA | 0.05512 | 0.01764 | 0.10149 | 0.0 | 463 | 766 |

Table 10: Version Information for Clinical Data Analysis

| Component | Version/Value |
|---|---|
| Code Hash | d57f11bc87fee69e |
| Timestamp | 2025-11-11 10:42:31 |
| Python Version | 3.13.3 |
| Pandas | 2.2.3 |
| NumPy | 2.2.5 |
| Scikit-learn | 1.7.2 |
| SciPy | 1.16.2 |
